# Synaptopathy in the TDP43ΔNLS Mouse Model of Sporadic Amyotrophic Lateral Sclerosis

**DOI:** 10.1101/2025.03.25.645173

**Authors:** Ani Ayvazian-Hancock, Emma Butler, Claire F. Meehan, Gareth B. Miles, Matthew J. Broadhead

## Abstract

Sporadic cases of Amyotrophic Lateral Sclerosis (sALS) represent the most common form of motor neuron disease. sALS is characterised by pathological cytoplasmic inclusions of TDP-43, so-called reactive astrocyte pathology, and motor neuron degeneration. Early-stage alterations in certain subpopulations of synapses between neurons are thought to be a key driver of the early pathological mechanisms of ALS. However, we do not have a clear understanding of which types of synapses are impacted in ALS. Identifying vulnerable synapses affected in sALS models may provide insights into the key sites of disease pathogenesis. In this study we have performed quantitative high-resolution microscopy to survey different synapse subtypes, including excitatory (glutamatergic), inhibitory (glycinergic) and modulatory (cholinergic C-Bouton) synapses, in the spinal cord of a mouse model of sALS showing inducible TDP-43 pathology (TDP43ΔNLS) restricted to neurons. We have identified changes in cholinergic synapses and a subpopulation of excitatory synapses. Mice display robust neuronal TDP-43 pathology and evidence of TDP-43 changes at cholinergic C-boutons. We also observe no evidence of astrocytic pathology nor changes in the fraction of synapses that are contacted by astrocytes, demonstrating that synapse pathology is driven by cell-autonomous (neuronal) mechanisms. Overall, our findings highlight the selective vulnerability of distinct synapse populations in ALS.

## Introduction

Amyotrophic Lateral Sclerosis (ALS) is a neurodegenerative disorder characterized by motor neuron (MN) cell death in the nervous system, which leads to behavioural symptoms including muscle weakening, and eventually death due to paralysis of respiratory muscles ^1,2^. ALS may be caused by a combination of genetic mutations or environmental risk factors ^3^. While a small proportion of ALS cases are inherited and classified as familial ALS (fALS), approximately 90% of patients have sporadic ALS (sALS), with no known family history of the disease ^4–6^. Over 30 different gene mutations are implicated in the aetiology of ALS, including mutations in genes encoding guanine nucleotide exchange chromosome 9 open reading frame 72 (C9ORF72), Cu/Zn superoxide dismutase 1 (SOD1) RNA-binding protein Fused in Sarcoma (FUS) and TAR DNA-binding protein 43 (TDP43).

Despite the complex and diverse genetic and molecular aetiology, approximately 97% of ALS cases display TDP-43 pathology, characterised by the mislocalisation and aggregation of hyper-phosphorylated TDP-43 inclusions in the cytoplasm of neurons ^7–11^. TDP-43 has many cellular functions including RNA spicing and translation ^12–14^. Whilst TDP-43 is predominantly expressed in the nucleus, it is also expressed in lower concentrations throughout the cell cytoplasm, including at presynaptic and postsynaptic compartments ^15,16^, where it may have a role in local translational processes related to synaptic proteins ^13,17–19^. TDP-43 pathology may represent a conserved mechanism driving neurodegeneration across the diverse spectrum of ALS patients.

Before the loss of MNs in the spinal cord, central chemical synapses between neurons are highly susceptible to structural and functional changes that may drive early-stage hyper-excitability in MNs, which is hypothesised to lead to excitotoxicity and subsequent MN cell death^20^. MNs are the final common pathway of the nervous system, receiving a wide range of different synaptic inputs. Synapses are highly diverse in their molecular composition, morphology, and function, both within and between different neural circuits and anatomical regions of the nervous system^21–25^. Synaptic diversity may also incur selective vulnerability of molecularly or structurally distinct synapse subtypes ^26,27^. For example, tripartite synapses have previously been shown to be selectively vulnerable to degeneration in ALS compared to synapses not contacted by astrocytes ^28^. A range of other studies indicate changes in other classes of synapses in ALS, including excitatory ^29,30^, inhibitory ^31–33^ and cholinergic synapses ^34–36^ in the spinal cord. Therefore, identifying which types of synapses are most significantly impacted in ALS is critical for understanding the mechanisms of the disease and may help identify critical pathways for targeted therapeutic strategies.

In this study, we examined different types of synapses in the lateral motor pools of the upper-lumbar spinal cord in a mouse model of sALS that drives TDP-43 mislocalisation and significant motor deficits ^36^. Using immunohistochemistry and Airyscan confocal microscopy, we identified selective vulnerability of certain synapse subtypes. Strikingly, however, we revealed a strong concordance in the hallmarks of synaptic pathology in subtypes of excitatory synapses between this TDP-43-associated model and other models harbouring different genetic mutations. These findings illustrate conserved hallmarks of synaptopathy across the diverse spectrum of ALS that may point towards a convergent mechanism.

## Methods

### Ethics

All procedures on live animals were approved by the Danish Animal Experiments Inspectorate (Permission number 2018-15-0201-01426) and were conducted in accordance with the EU Directive 2010/63/EU. Once mouse tissue was shipped to the UK, all procedures were conducted in accordance with the Scientific Procedures UK Animals Act 1986.

### TDP43ΔNLS Mouse Model & Behavioural Assessment

Mice were produced by crossing a tetO-hTDP-43-ΔNLS line 4 (https://www.jax.org/strain/014650) with a NEFH-tTA line 8 (https://www.jax.org/strain/025397). This cross breeding created a mouse model with tetracycline-repressible TDP-43 which lacks the nuclear localization signal. When the tetracycline analogue, Doxycycline, is removed from the diet of bigenic animals, toxic mislocalisation of TDP-43 into the cytoplasm of neurons is induced ^37^. Mice were genotyped using a standard PCR assay described in previous study ^36^. Four transgene-specific primers were used for PCR amplification to enable the final detection of these sequences on electrophoretic gels; 5′-CTC GCG CAC CTG CTG AAT-3′ (Tg(NEFH-tTA) forward), 5′-CAG TAC AGG GTA GGC TGC TC-3′(Tg(NEFH-tTA) reverse), 5′-TTG CGT GAC TCT TTA GTA TTG GTT TGA TGA-3′ (Tg(tetO-hTDP-43-ΔNLS) forward) and 5′-CTC ATC CAT TGC TGC TGC G-3′ (Tg(tetO-hTDP-43-ΔNLS) reverse) ^36,38^. In our study, all mice underwent withdrawal of doxycycline from the diet at the same time point. Mice expressing only the tTA gene were designated as Control animals, whilst mice expressing both the NEFH-tTA and TetO promoter (bigenic) were designated as TDP43ΔNLS animals exhibiting symptoms of sALS. The ALS-like phenotype was confirmed using simple behavioural test. In the two-minute tail suspension test, mice are gently suspended by their tail for up to two minutes. Healthy mice will normally extend out their hind limbs during this task but mice with motor dysfunction display a distinctive hindlimb clasping behaviour. In the grip strength endurance test, mice are placed on a metal grid which is gently shaken to encourage them to grip on. it is then inverted over a soft surface and the endurance time that the mouse can hold on for is recorded (up to two minutes). The mean of three trials (with a ten minute break between each trial) was calculated.

### Tissue Collection

Animals were euthanized and tissue was collected at 11 weeks of age (4 weeks post-induction). Euthanasia was performed with Sodium Pentobarbital (120 mg/kg), and animals transcardially perfused with 0.9% saline followed by a 4% paraformaldehyde. Spinal cords were then removed and post-fixed for 2 hours in 4% paraformaldehyde. Spinal cords were then immersed in 30% sucrose in phosphate buffered saline (PBS) overnight for cryoprotection. Tissue was then cryo-embedded in OCT compound and stored at −80°C. Cryosections, at 20 μm thickness, were obtained using a Leica CM1860 cryostat, mounted on Leica X-tra adhesive slides and stored at -80°C for long term preservation.

### Immunohistochemistry

Immunohistochemistry for neurons, synapses and astrocytes was conducted as described previously ^16,24,28,39^. Slides with tissues sections were thawed in a benchtop heater at 37°C for 30 min to aid the adherence of the tissue to the glass slides and reduce tissue loss during subsequent wash steps. Slides were washed three times in PBS. Hydrophobic pens were used to draw rings around each spinal cord tissue on the slides, and the tissue was then incubated in PBS containing 3% Bovine Serum Albumin (BSA; Sigma Aldrich) and 0.2% Triton X100 (Fischer,) for 2 h at room temperature to block non-specific binding and permeabilise the tissue. Primary antibodies were diluted 1:500 in PBS containing 1.5% BSA and 0.1% Triton X100, and samples were incubated with primary antibody solution for 2 nights at 4°C. Slides were then washed in 1X PBS five times over the course of one hour. Secondary antibodies were then added to slides, diluted 1:500 in 0.1% Triton X, and incubated for 1.5 hours. Slides were washed an additional five times in 1X PBS over the course of an hour. If nuclear labelling was performed, DAPI stain was diluted in deionized (DI) water at 1:4000 dilution and applied to the slides for 10 minutes before being washed in DI water to stop the reaction. Finally, the slides were then dried, and 1.5mm thick cover slips were applied with Prolong Glass Antifade Mountant (Invitrogen, P36980).

Primary and secondary antibodies used are detailed in Table 1.

| Primary Antibody Species and Target | Primary Antibody Source | Secondary Antibody Species & Fluorophore | Secondary Antibody Source |
| --- | --- | --- | --- |
| **Mouse anti-VGLUT2** | Abcam<br>ab79157 | Donkey anti-Mouse Alexa<br>Fluor 405 | Invitrogen<br>A31573 |
| **Guinea Pig anti-VGLUT1** | Millipore<br>AB5905 | Donkey anti-Guinea Pig Alexa<br>Fluor 647 | Jackson Laboratories<br>706-605-148 |
| **Rabbit anti-pTDP43** | ProteinTech<br>22309-01 | Donkey anti-Rabbit Alexa<br>Fluor 555 | Invitrogen,<br>A31570 |
| **Camelid anti-PSD95** | Synaptic Systems N3702-At488-L | Not Applicable | Not applicable |
| **Guinea Pig anti-GLYT2** | Synaptic Systems<br>272-004 | Donkey anti-Guinea Pig Alexa<br>Fluor 647 | Jackson Laboratories<br>706-605-148 |
| **Goat anti-VACHT** | Millipore<br>ABN100 | Donkey anti-Goat Alexa Fluor<br>488 | Invitrogen,<br>A11055 |
| **Goat anti-MMP9** | Sigma<br>M9570 | Donkey anti Goat Alexa Fluor<br>488 | Invitrogen,<br>A11055 |
| **Mouse anti-SMI32** | Biolegend<br>801701 | Donkey anti-mouse Fluor647 | Abcam<br>Ab150107 |
| **Rabbit anti-pEzrin** | Abcam<br>ab47293 | Donkey anti-Rabbit Alexa<br>Fluor 555 | Invitrogen,<br>A31570 |
| **Rabbit anti-GFAP** | DAKO<br>Z0334 | Donkey anti-Rabbit Alexa<br>Fluor 555 | Invitrogen<br>A31570 |
| **Chicken anti-GFAP** | Invitrogen<br>PA110004 | Donkey anti-Chicken Alexa<br>Fluor 594 | Sigma<br>SAB4600094 |

### Airyscan Confocal Microscopy

High-resolution imaging of synapses was performed using the Zeiss LSM800 laser scanning confocal microscope, based on an Axio ‘Observer 7’ microscope, equipped with a 63× 1.4 NA objective lens, an Airyscan super-resolution module plus two individual GaAsP PMT detectors. Illumination was provided by 405, 509, 561, and 640 nm laser lines. The pixel size was set to 0.04 μm, resulting in an image size of 78 x 78 µm (2210 x 2210 pixels). Single-z plane images were acquired with bidirectional scanning and 2x averaging, with a pixel scanning time of 2.4 μs. Airyscan processing on images was performed to yield sub-diffraction limit resolution images of structures, with the Wiener filter kept the same for images within a data set. Images were acquired in the lateral ventral horn, approximately lamina IV, containing pools of MNs.

### Andor BC43 Confocal Microscopy

Low-magnification high-speed confocal images of mouse lumbar spinal cord tissue were acquired with an Andor BC43 benchtop confocal equipped with 405, 488, 561 and 638 nm lasers for illumination. Z-stack images were acquired at 20× 0.8 NA objective lens. Each multi-channel acquisition consisted of 11 images acquired across a 2 µm range in Z-axis with resultant voxel size of 311 x 311 x 181 nm.

### Image analysis

Image analysis was conducted in FIJI ^40^. Researchers were blinded to the genotype of animals to prevent bias. Image processing and analysis of synaptic, astrocytic and pTDP-43 structures was performed using customised macros written in FIJI as used and reported in previous studies ^16,24,28^. Briefly, images undergo processing steps including background subtraction and Gaussian smoothing before being binarized by thresholding. Watershed splitting was applied to segment structures that became merged following thresholding. Small structures consisting of less than 8 pixels were filtered out. From the processed images, the number, size and fluorescence intensity of particles was quantified by redirecting the analysis of the binarized images back to the original raw image. Structures were deemed to colocalize when particles overlapped by a minimum of 1 pixel. Analysis of pTDP-43 expression in cell cytoplasm and nuclei was performed by manually delineating nuclei guided by DAPI labelling and the surrounding soma (excluding the nuclei) guided by the expression of pTDP-43 itself. Analysis was restricted to large cells with robust pTDP-43 expression in the lateral motor pools of the lumbar spinal cord and were thus predominantly neurons. Morphological analysis of astrocytes using GFAP labelling was performed by binarizing GFAP images after background subtraction and mean smoothing to remove noise, followed by skeletonising the binarized image and using the AnalyzeSkeleton plugin.

### Data Analysis

Data was collated and managed in Microsoft Excel. Statistical analysis was performed in SPSS (IBM, version 28.0). Measurements of synapse or pTDP-43 cluster density (number per unit area) and size were averaged (median) across all images acquired from at least two tissue sections from individual mice (n). Shapiro Wilks test for normality was performed for measures of synapse size and number, with all data showing normal distributions (SPSS 25.0, IBM). Statistical analysis was performed using a Two-Sample T Test. Graphs were generated in Microsoft Excel plotting the mean ± standard error of the mean and figures were constructed in Powerpoint.

## Results

### Motor Phenotype in the Mouse Model of sALS

Behavioural test confirmed that the mice used for this investigation display the same motor phenotypes as has previously been described for this model ^36–38^. All TDP43ΔNLS mice showed hindlimb clasping within 5 seconds of being suspended by their tail. None of the control mice exhibited clasping behaviour during this test. The endurance time on the grip strength endurance test was significantly reduced in the TDP43ΔNLS mice, whereas all control mice could complete the task for the full 2 minutes (Figure 1).

**Figure 1.**
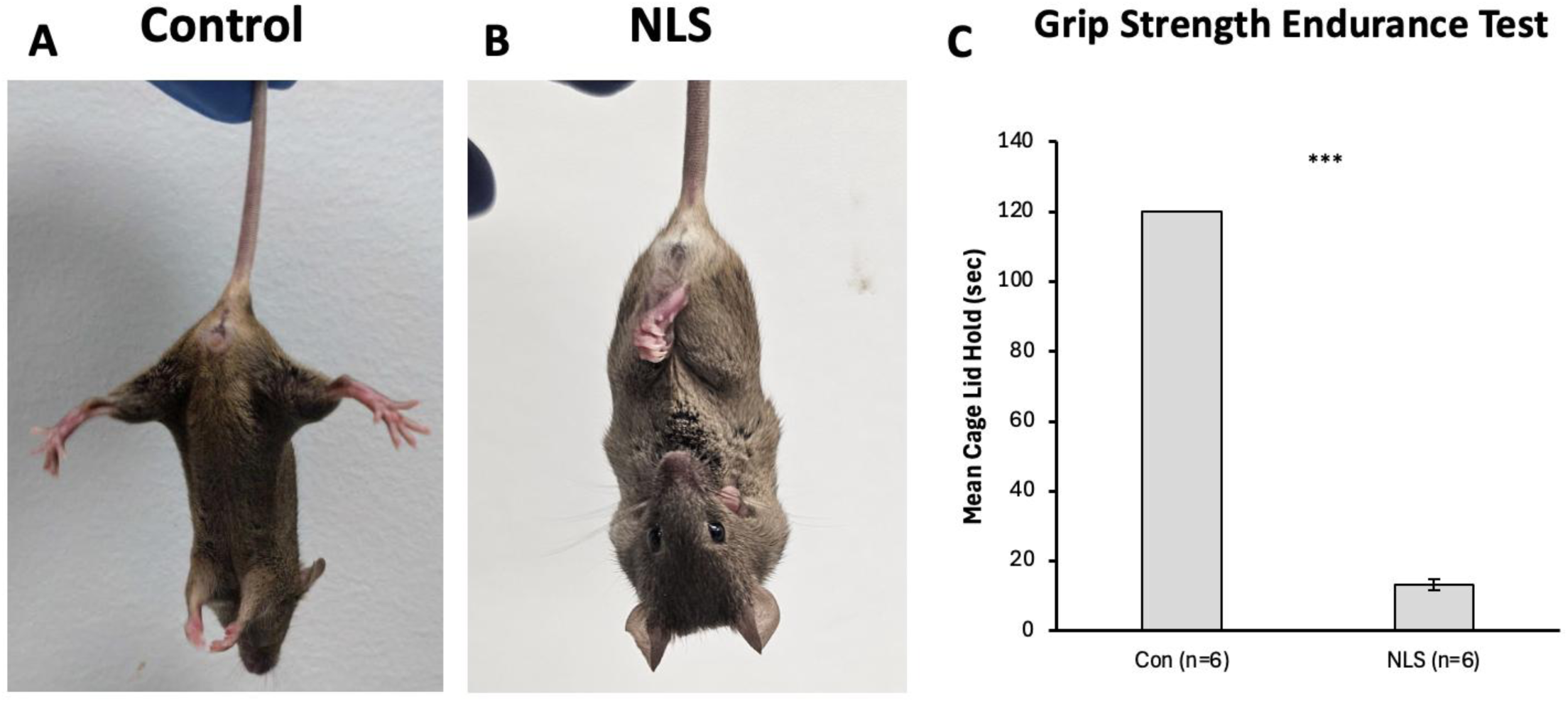
Behavioral images and quantification of mouse phenotype. (A) Image of non-transgenic control mouse displaying a healthy splayed limb phenotype. (B) Image of TDP43ΔNLS mouse model displaying a severe clasping phenotype affecting all 4 limbs. (C) Bar graph representation of grip strength endurance test showed a significant decrease in mean cage lid holding time in the TDP43ΔNLS mouse model compared to non-transgenic controls (t(10)=71.499, p<0.001).

### Cellular Pathology in Mouse Model of sALS

The main investigation began by characterising cellular pathological hallmarks in the spinal cord tissue of the TDP43ΔNLS model. To do this, we investigated TDP-43 pathology, MN pathology and astrocyte reactivity.

Firstly, immunohistochemistry and high-resolution microscopy was performed on tissue from our experimental cohort of control and TDP43ΔNLS mice, to visualise the expression of pTDP-43 in cells with DAPI labelled nuclei in the ventral horn of the mid-lumbar spinal cord. pTDP-43 labelling was predominantly localised to the nucleus, although punctate clusters of TDP-43 were also seen throughout cell bodies (Figure 2A). To verify the expected mislocalisation of pTDP-43 in mutant animals, the ratio of pTDP-43 within DAPI-labelled nuclei to pTDP-43 in the surrounding cell body was quantified. In TDP43ΔNLS spinal tissue, cells displayed a reduction or absence of nuclear pTDP-43 labelling compared to controls (t(10)=4.343, p=0.002) (Figure 2B). This confirmed that the experimental cohort of TDP43ΔNLS mice used in this study to model sALS displayed the expected TDP-43 pathology.

**Figure 2.**
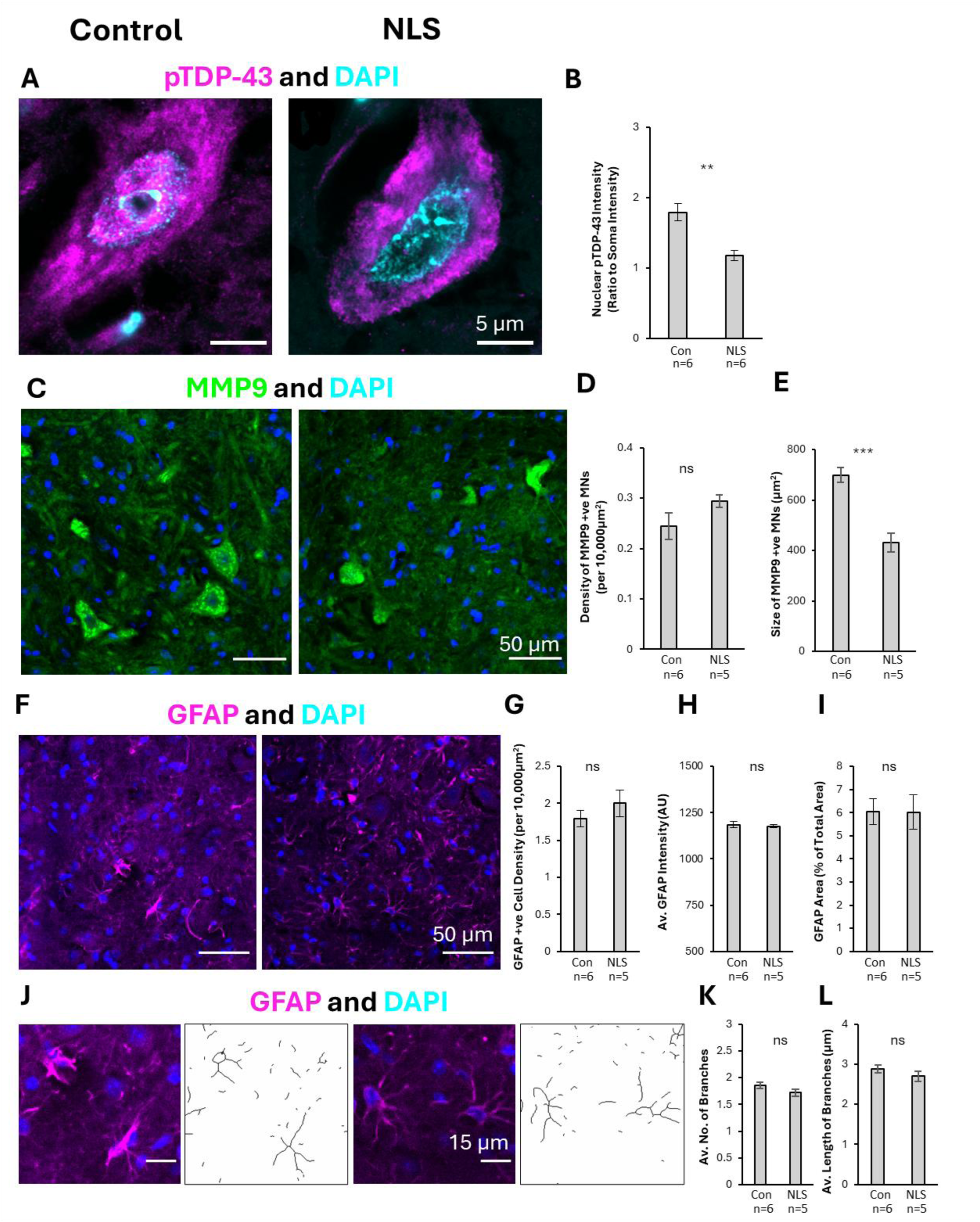
Characterising molecular and cellular pathology in TDP43ΔNLS mouse model. (A) Immunohistochemical images staining for pTDP-43 and DAPI in order to quantify the localization of pTDP-43 to the nucleus in both the TDP43ΔNLS mouse model and controls. (B) Bar graph representation of nuclear localization of pTDP-43 found significantly less nuclear pTDP-43 in the TDP43ΔNLS mouse model compared to non-transgenic controls (t(10)=4.34, p=0.002). (C). Immunohistochemical image assessment of motor neurons using MMP9+ ve and DPAI stains in order to quantify the density and size of motor neurons in the ventral horn of the lumbar spinal cord. (D) Bar graph representation of density of motor neurons showed no significant difference between the TDP43ΔNLS mouse model and non-transgenic controls (t(9)=1.351, p=0.201). (E) Bar graph representation of size of motor neurons showed a significant reduction in the size of motor neurons in the TDP43ΔNLS mouse model when compared to controls (t(9)=5.760, p<0.001). (F) Immunohistochemical assessment of astrocytes in the TDP43ΔNLS mouse model utilizing a GFAP and DAPI stain. (G) Bar graph assessment of GFAP cell density in the ventral horn of the mouse lumbar spinal cord, which showed no significant changes between the ΔNLS mouse model and controls (t(9)=1.195, p=0.262). (H) Bar graph assessment of GFAP intensity showed no significant changes between the experimental condition and controls (t(9)=0.427, p=0.679). (I). Bar graph assessment of total GFAP area showed no significant changes between ΔNLS mouse model and non-transgenic controls (t(9)=0.107, p=0.917). (J) Immunohistochemical images and masks of GFAP to determine number and length of astrocytic branches in ΔNLS mouse model compared to controls. (K) Bar graph assessment of number of astrocytic branches showed no significant difference between ΔNLS mice when compared with controls (t(9)=1.710, p=0.121). (L) Bar graph assessment of length of astrocytic branches revealed no significant changes between the ΔNLS mouse model and controls (t(9)=1.151, p=0.279).

We next asked whether the TDP43ΔNLS mice displayed MN pathology or reactive gliosis. Spinal cord sections were immunolabelled for MMP9 to label a subset of large alpha-MNs that innervate fast-twitch muscles (so-called fast motor neurons), GFAP to label astrocytes, and DAPI to identify cell nuclei. Analysis was conducted to assess changes in the number and size of MMP9-positive alpha-MNs in the TDP43ΔNLS mice compared to controls (Figure 2C). There was no difference in the density of motor neurons in the ventral horn of the lumbar spinal cord, as quantified from the number of MMP9 neurons per 10,000 µm^2^ (t(9)=1.351, p=0.201) (Figure 2D). There was, however, a significant reduction in the size of these motor neurons in the ΔNLS mice, when compared with controls (t(9)=5.760, p<0.001) (Figure 2E). This indicates that existing MNs are significantly impacted by the induced TDP-43 pathology in this model, though the phenotype does not yet lead to significant loss of MNs.

Next, GFAP-labelling was analysed to determine whether the TDP43ΔNLS mice displayed any signs of reactive gliosis. A range of parameters were measured, including the density of astrocytes, fluorescence intensity of GFAP, the overall area covered by GFAP labelling, and the branching morphology of the astrocytes (Figure 2F & 2J). Analysis revealed no changes in the density of the GFAP stained cells (t(9)=1.195, p=0.262) (Figure 2G), the fluorescence intensity of GFAP labelling (t(9)=0.427, p=0.679) (Figure 2H), or the fraction of area covered by GFAP labelling (t(9)=0.107, p=0.917) (Figure 2I) between the TDP43ΔNLS mouse model and non-transgenic controls. Furthermore, there were no significant differences in the number of branch points (t(9)=1.710, p=0.121) (Figure 2K) or the length of branch points(t(9)=1.151, p=0.279) of GFAP-positive astrocytes between the TDP43ΔNLS mice and controls (Figure 2L). This suggests that astrocytes may not be affected by the TDP43ΔNLS mutation present in this model.

In summary, this characterisation reveals that the TDP43ΔNLS mice used in this study displayed TDP-43 pathology and morphological changes in MNs in the spinal cord which is consistent with human ALS pathology ^9^ and previous findings using this mouse model ^36^. However, unlike what is seen in human ALS pathology ^41^, the TDP43ΔNLS mouse model does not show overt astrocytic reactivity.

### Differential Vulnerability of Synapses in sALS Model

We next performed a series of immunohistochemical experiments to measure changes in synapse marker expression in TDP43ΔNLS mice. Cholinergic C-bouton synapses were visualized with vesicular acetylcholine transporter (VAChT) labelling (Figure 3Ai). Glycine transporter 2 (GLYT2) labelling was performed to identify presynaptic boutons of inhibitory synapses (Figure 3Bi). Two different types of excitatory synapses were visualised by labelling for the vesicular glutamate transporter 1 (VGLUT1) (predominantly proprioceptive inputs) (Figure 3Ci) and 2 (VGLUT2) (predominantly synapses from excitatory spinal interneurons) (Figure 3Di). The expression of these four different presynaptic markers was analysed to measure changes in their density (number of synapses per unit area), total synaptic coverage (sum of areas of synaptic labelling) and synapse size (area in μm^2^).

**Figure 3.**
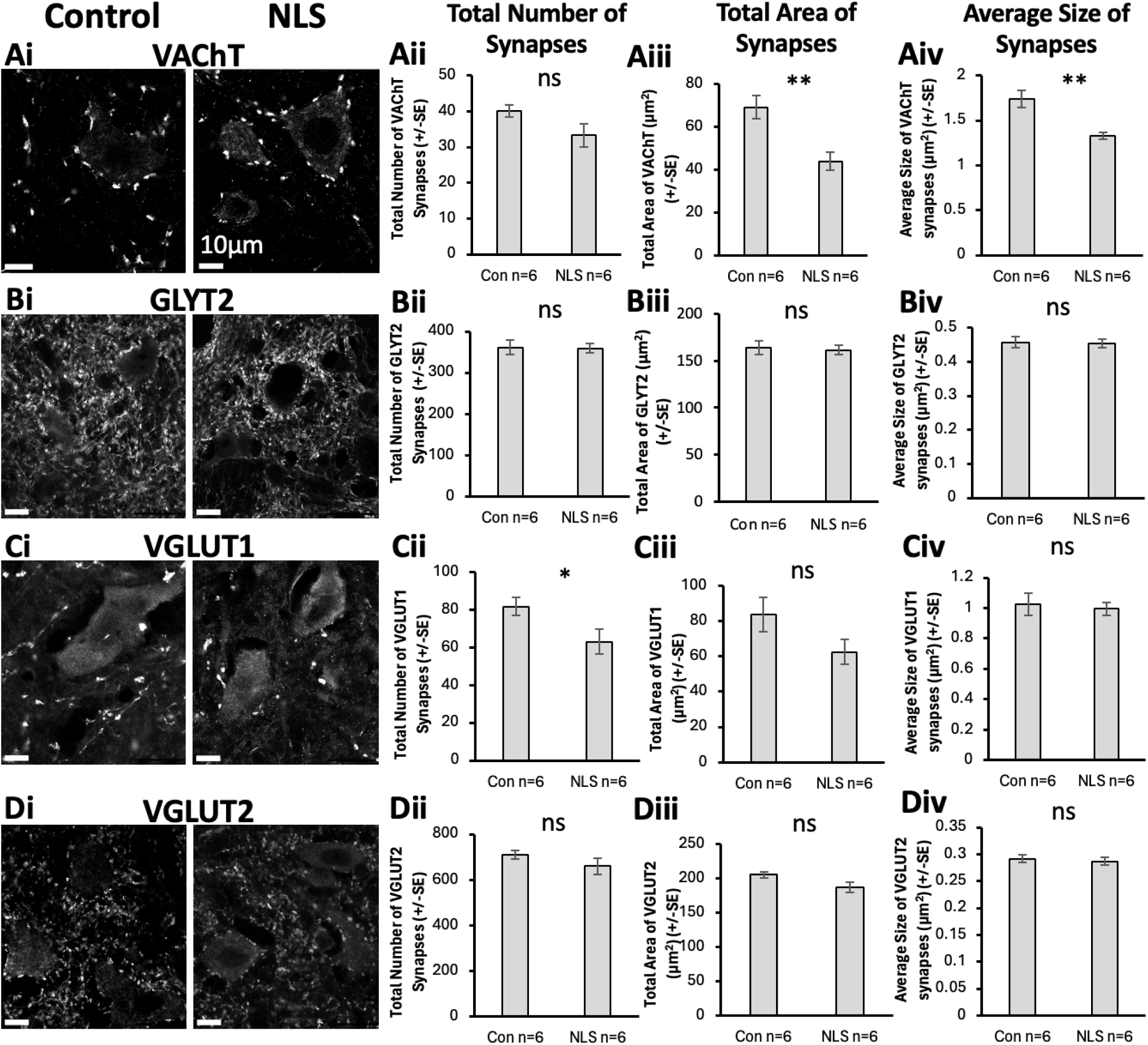
VAChT, GLYT2, VGUT1, and VGLUT2 general synaptic assessment. (Ai) Immunohistochemical staining of VAChT synapses. (Aii-Aiv) Bar graph representation of total number, area, and size of VAChT synapses showing no significant changes in the number of VAChT synapses (t(10)=1.848, p=0.094), but a significant reduction in the total area coverage (t(10)=3.805, p=0.003) and average size of VAChT synapses (t(10)=3.932, p=0.003) in the TDP43ΔNLS mice compared to controls. (Bi) Immunohistochemical staining of GLYT2 synapses. (Bii-Biv) Bar graph representation of total number (t(10)=0.104, p=0.919), area (t(10)=0.265, p=0.796), and size (t(10)=0.146, p=0.444) of GLYT2 synapses showing no significant changes between TDP43ΔNLS mice and controls. (Ci) Immunohistochemical staining of VGLUT1 synapses. (Cii-Civ) Bar graph representation of total number, area, and size of VGLUT1 synapses showed a significant reduction in the number of VLGUT1 synapses in the TDP43ΔNLS mouse model compared to controls (t(10)=2.278, p=0.046), but no differences in the total area coverage (t(10)=1.793, p=0.103) or average size (t(10)=0.364, p=0.723) of VGLUT1 synapses between conditions. (Di) Immunohistochemical staining of VGLUT2 synapses. (Dii-Div) Bar graph representation of total number (t(10)=1.259, p=0.237), area (t(10)=2.059, p=0.066), and size (t(10)=0.417, p=0.685) of GLYT2 synapses showing no significant changes between TDP43ΔNLS mice and controls.

Analysis of VAChT synapses revealed that there was no significant difference in their density, despite an overall trend towards fewer VAChT-boutons in the TDP43ΔNLS mice compared to controls (t(10)=1.848, p=0.094) (Figure 3Aii). However, the total area covered by VAChT labelling was significantly less in TDP43ΔNLS mice compared to controls (t(10)=3.805, p=0.003) (Figure 3Aiii). Similarly, the size of VAChT boutons was significantly smaller in TDP43ΔNLS mice compared to controls (t(10)=3.932, p=0.003) (Figure 3Aiv).

Analysis of GLYT2-labelling revealed that there was no significant difference in the number of inhibitory synapses between TDP43ΔNLS mice and controls (t(10)=0.104, p=0.919) (Figure 3Bii). Similarly, there was no difference in the total area covered by GLYT2 labelling (t(10)=0.265, p=0.796) (Figure 3Biii), or the size of inhibitory presynaptic boutons (t(10)=0.146, p=0.444) (Figure 3Biv), in the TDP43ΔNLS mice compared to controls.

Excitatory synapses were visualized using both VGLUT1 (Figure 3Ci) and VGLUT2 (Figure 3Di) labelling. We found a lower density of VGLUT1 synapses in the TDP43ΔNLS mice compared to controls (t(10)=2.278, p=0.046) (Figure 3Cii). Meanwhile, there was no difference between TDP43ΔNLS mice and controls in the total area of VGLUT1 labelling (t(10)=1.793, p=0.103) (Figure 3Ciii) or the average size of VGLUT1 boutons (t(10)=0.364, p=0.723) (Figure 3Civ). In contrast, there was no difference in VGLUT2 synaptic density (t(10)=1.259, p=0.237) (Figure 3Dii), total area covered by VGLUT2 labelling (t(10)=2.059, p=0.066) (Figure 3Diii), or the size of VGLUT2 boutons (t(10)=0.417, p=0.685) (Figure 3Div) in TDP43ΔNLS mice compared to controls.

From analysis of these individual synaptic markers in areas around MNs in the mouse spinal cord, significant and selective synaptic pathology is observed. Our results highlight that cholinergic C-boutons and VGLUT1-associated excitatory synapses are most significantly impacted in the TDP43ΔNLS model of sALS, whilst inhibitory synapses (GLYT2) and the majority of excitatory synapses (VGLUT2) are not significantly affected.

### TDP-43 Expression in Different Synapse Subtypes

Following the assessment of cholinergic, inhibitory, and excitatory synapses (Figure 3), and the previous observation of cytoplasmic mis-localisation of TDP-43 (Figure 2A-B), the presence of pTDP-43 pathology within these synapses was examined.

We found a significantly lower frequency of VAChT boutons that contained pTDP-43 (approximately 74%) in TDP43ΔNLS mice compared to non-transgenic controls (approximately 80%) (t(10)=2.405, p=0.037) (Figure 4Aii). Additionally, there was significant fewer pTDP-43 clusters per VAChT bouton in the TDP43ΔNLS model (approximately 2 clusters per bouton) compared to controls (approximately 3 clusters per bouton) (t(10)=3.946, p=0.003) (Figure 4Aiii). In examining the size of the pTDP-43 clusters localized within the VAChT boutons, there was no significant difference (t(10)=2.020, p=0.071) (Figure 4Aiv).

**Figure 4.**
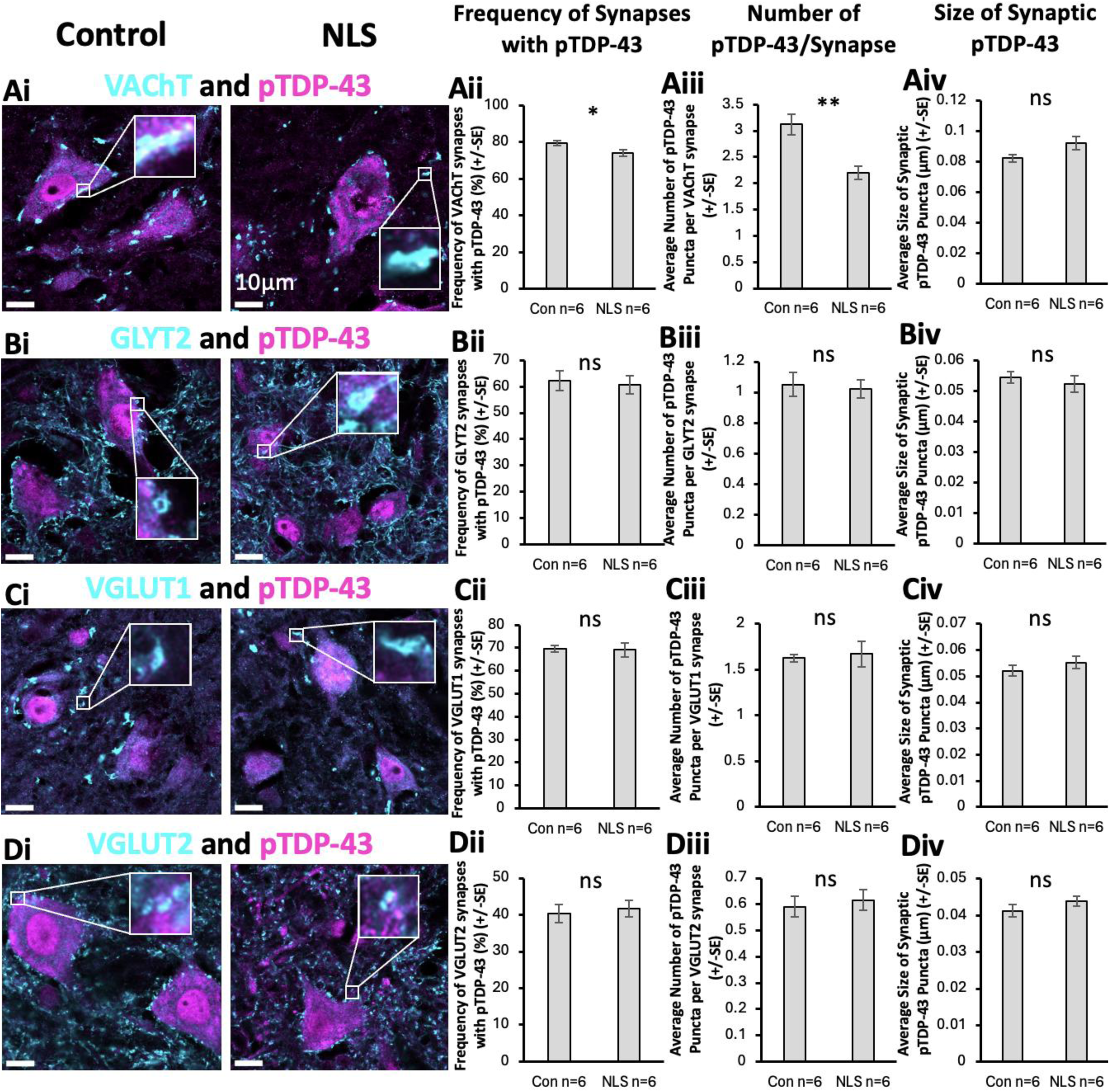
Assessment of pTDP-43 associated with VAChT, GLYT2, VGLUT1, and VGLUT2 synapses. (Ai) Immunohistochemical stain of VAChT and pTDP-43 to assess synaptically colocalized pTDP-43 expression. (Aii-Aiv) Bar graph representation of pTDP-43 colocalized to VAChT showing a significantly reduced frequency of VAChT synapses colocalized to VAChT (t(10)=2.405, p=0.037), a significantly reduced number of pTDP-43 puncta colocalized per VAChT synapse (t(10)=3.946, p=0.003), and no change in the size of pTDP-43 colocalized to VAChT synapses (t(10)=2.020, p=0.071), in TDP43ΔNLS mice compared to controls. (Bi) Immunohistochemical stain of GLYT2 and pTDP-43 to assess synaptically colocalized pTDP-43 expression. (Bii-Biv) Bar graph representations showing no changes in frequency (t(10)=0.318, p=0.757), puncta per synapse (t(10)=0.280, p=0.785), or size of pTDP-43 colocalized with GLYT2 synapses (t(10)=0.651, p=0.529), between experimental conditions. (Ci) Immunohistochemical stain of GLYT2 and pTDP-43 to assess synaptically colocalized pTDP-43 expression. (Cii-Civ) Bar graph representations showing no changes in frequency (t(10)=0.147, p=0.886), puncta per synapse (t(10)=0.320, p=0.756), or size of pTDP-43 colocalized with VGLUT1 synapses (t(10)=0.966, p=0.357), between experimental conditions. (Di) Immunohistochemical stain of GLYT2 and pTDP-43 to assess synaptically colocalized pTDP-43 expression. (Dii-Div) Bar graph representations showing no changes in frequency (t(10)=0.398, p=0.699), puncta per synapse (t(10)=0.448, p=0.664), or size of pTDP-43 colocalized with VGLUT2 synapses (t(10)=1.225, p=0.249), between experimental conditions

No significant differences were observed in the frequency of GLYT2 synapses containing pTDP-43 (t(10)=0.318, p=0.757) (Figure 4Bii), the number of pTDP-43 puncta colocalized per GLYT2 synapse (t(10)=0.280, p=0.785) (Figure 4Biii), or the average size of the pTDP-43 puncta associated with GLYT2 synapses (t(10)=0.651, p=0.529) (Figure 4Biv), when comparing the TDP43ΔNLS mouse model and non-transgenic controls.

Similarly, no differences were seen between groups when assessing the frequency of pTDP-43 colocalized with VGLUT1 synapses (t(10)=0.147, p=0.886) (Figure 4Cii), the number of pTDP-43 puncta per VGLUT1 synapse (t(10)=0.320, p=0.756) (Figure 4Ciii), or the size of pTDP-43 associated with VGLUT1 synapses (t(10)=0.966, p=0.357) (Figure 4Civ). Finally, there was also no difference in the frequency of VGLUT2 synapses associated with pTDP-43 (t(10)=0.398, p=0.699) (Figure 4Dii), the number of pTDP-43 puncta colocalized per VGLUT2 synapse (t(10)=0.448, p=0.664) (Figure 4Diii), or the size of the pTDP-43 puncta associated with VGLUT2 synapses (t(10)=1.225, p=0.249) (Figure 4Div).

Our findings show synapse subtype-specific changes in TDP-43 expression. We observed reduced association of TDP-43 with C-boutons, however, the other excitatory and inhibitory synapse populations showed no evidence of changes in their content of TDP-43.

### Tripartite Synapses in sALS model

We have previously shown that excitatory tripartite synapses are selectively vulnerable to degeneration in the commonly used SOD1^G93a^ model of fALS, and in human cases based on analysis of post-mortem tissue from ALS patients with C9ORF72 mutations ^28^. We therefore hypothesised that tripartite synapses may also represent selectively vulnerable synapse subtypes in the TDP43ΔNLS mouse model of sALS ^28^

Excitatory synapses were examined by labelling for presynaptic VGLUT1 and VGLUT2 and the postsynaptic marker PSD95. For these assessments, synapses were defined as presynaptic boutons partially colocalised with PSD95. Synapses were assigned as tripartite or non-tripartite based on partial colocalization with the perisynaptic astrocytic process (PAP) marker, phosphorylated-Ezrin (p-Ezrin) Approximately 75% of VGLUT1-PSD95 synapses in the control animals were classified as tripartite synapses based on their association with p-Ezrin puncta (Figure 5A). There was no significant difference in the percentage of VGLUT1-synapses that were tripartite between TDP43ΔNLS and control mice (t(10)=0.243, p=0.813) (Figure 5B). The size of VGLUT1 boutons from all bona fide synapses was no different between TDP43ΔNLS and control mice (t(10)=0.571, p=0.580) (Figure 5C). Similarly, there was no difference in VGLUT1 bouton size between TDP43ΔNLS and control mice whether they were part of tripartite synapses (t(10)=0.511, p=0.620) or non-tripartite synapses (t(10)=1.230, p=0.247) (Figure 5C). Whilst there was no change in VGLUT1 presynaptic bouton morphology, the size of the opposed PSD95 puncta (the PSDs) was significantly smaller in TDP43ΔNLS mice compared to controls (t(10)=4.014, p=0.00246). When synapses were analysed based on their association with p-Ezrin PAPs, the PSDs that were part of non-tripartite synapses were significantly smaller in TDP43ΔNLS mice compared to controls (t(10)=2.946, p=0.015). In contrast, PSDs that were part of tripartite synapses were not significantly smaller in TDP43ΔNLS mice (∼0.19 µm^2^) compared to controls (∼0.17 µm^2^) (Figure 5D).

**Figure 5.**
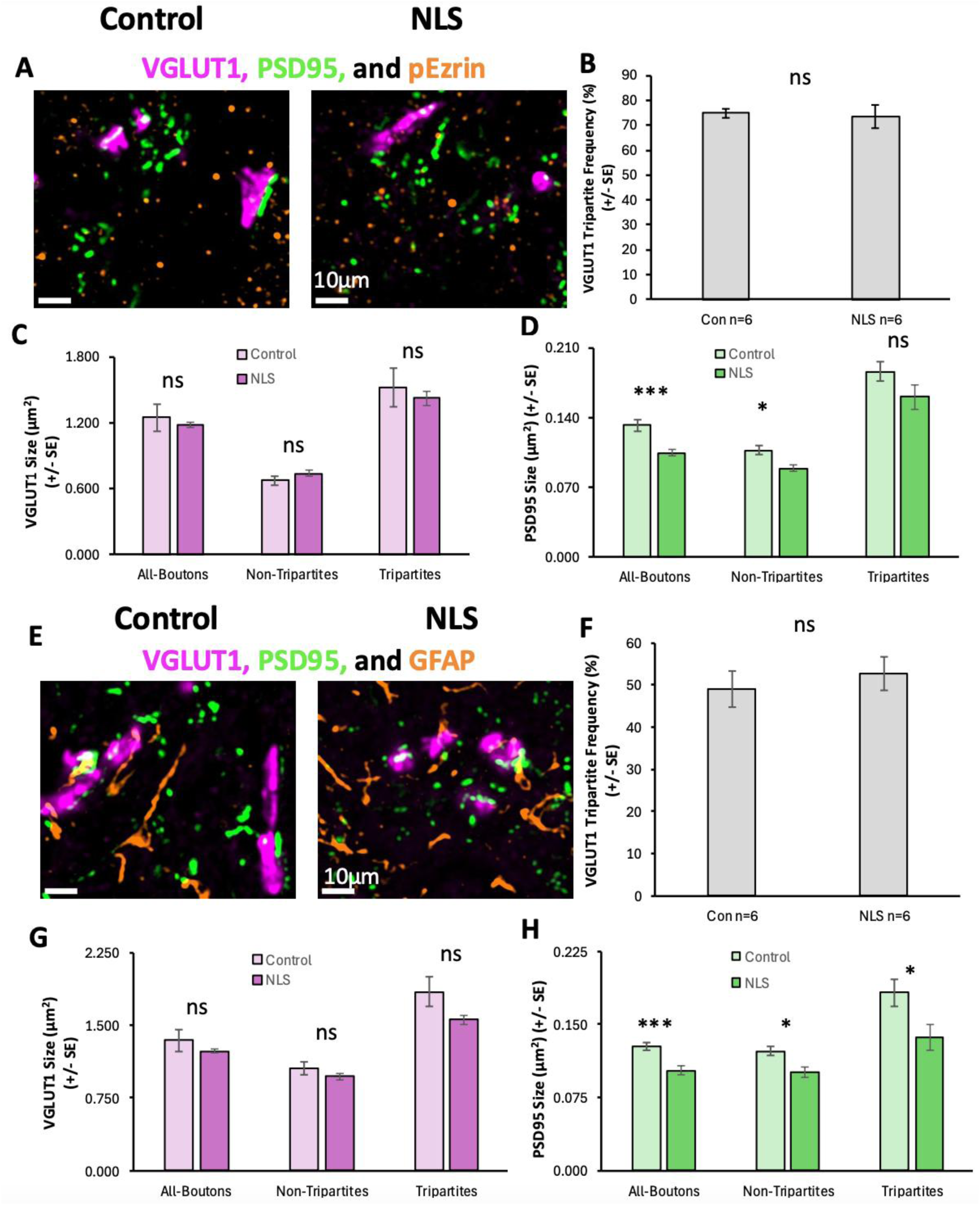
Excitatory VGLUT1-associated Tripartite Synapses in sALS. (A) Immunohistochemical stain of VGLUT1, PSD95 and pEzrin to identify excitatory tripartite synapses. (B) Bar graph plotting the percentage of VGLUT1 synapses that are tripartite (pEzrin contacted) in control and TDP43ΔNLS (NLS) mice. (C) Bar graph of the size of VGLUT1 boutons associated with all synapses, tripartite and non-tripartite synapses, in control and NLS mice. (D) Bar graph of the size of PSDs associated with all VGLUT1 synapses, tripartite and non-tripartite synapses in control and NLS mice. (E) Immunohistochemical stain of VGLUT1, PSD95 and GFAP to identify excitatory tripartite synapses. (F) Bar graph plotting the percentage of VGLUT1 synapses that are tripartite (GFAP contacted) in control and TDP43ΔNLS (NLS) mice. (G) Bar graph of the size of VGLUT1 boutons associated with all synapses, tripartite and non-tripartite synapses, in control and NLS mice. (H) Bar graph of the size of PSDs associated with all VGLUT1 synapses, tripartite and non-tripartite synapses in control and NLS mice

For further validation, we repeated this experiment with an alternative astrocytic marker, GFAP (Figure 5E). There was also no difference in the percentage of VGLUT1 synapses contacted by pEzrin (approximately 50%) between control and TDP43ΔNLS mice (Figure 5F). Similarly, there was no difference in the size of VGLUT1 presynaptic boutons, whether they were tripartite synapses (T(10)=1.77, p=0.108) or non-tripartite synapses (T(10)=1.14, p=0.282) (Figure 5G). We observed that the PSDs associated with all VGLUT1 synapses were significantly smaller in TDP43ΔNLS mice compared to controls (t(10)=3.666, p=0.004), whether they were associated with tripartite synapses (t(10)=2.413, p=0.037) or non-tripartite synapses (t(10)=3.047, p=0.0123) (Figure 5H). Taken together, our data indicate that VGLUT1 synapses display postsynaptic alterations that are independent of astrocytic interactions.

Analysis of VGLUT2-associated synapses and p-Ezrin-associated tripartite synapses revealed that approximately 38% of all VGLUT2-associated excitatory synapses were classified as tripartite synapses based on their association with p-Ezrin puncta (Figure 6A). There was no significant difference in the percentage of VGLUT2-synapses that were tripartite between TDP43ΔNLS and control mice (t(10)=0.014, p=0.989) (Figure 6B). The size of VGLUT2 presynaptic boutons was not significantly different between TDP43ΔNLS and control mice (t(10)=1.036, p=0.325) (Figure 6C). There was also no difference in VGLUT2 bouton size between TDP43ΔNLS and control mice whether they were part of tripartite synapses (t(10)=0.108, p=0.916) or non-tripartite synapses (t(10)=0.648, p=0.532) (Figure 6C). When we analysed the PSDs of VGLUT2-associated synapses, there was also no significant difference in their size between TDP43ΔNLS mice and controls, whether from all VGLUT2-associated synapses (t(10)=0.708, p=0.495), tripartite synapses (t(10)=0.117, p=0.910) or non-tripartite synapses (t(10)=0.686, p=0.508) (Figure 6D).

**Figure 6.**
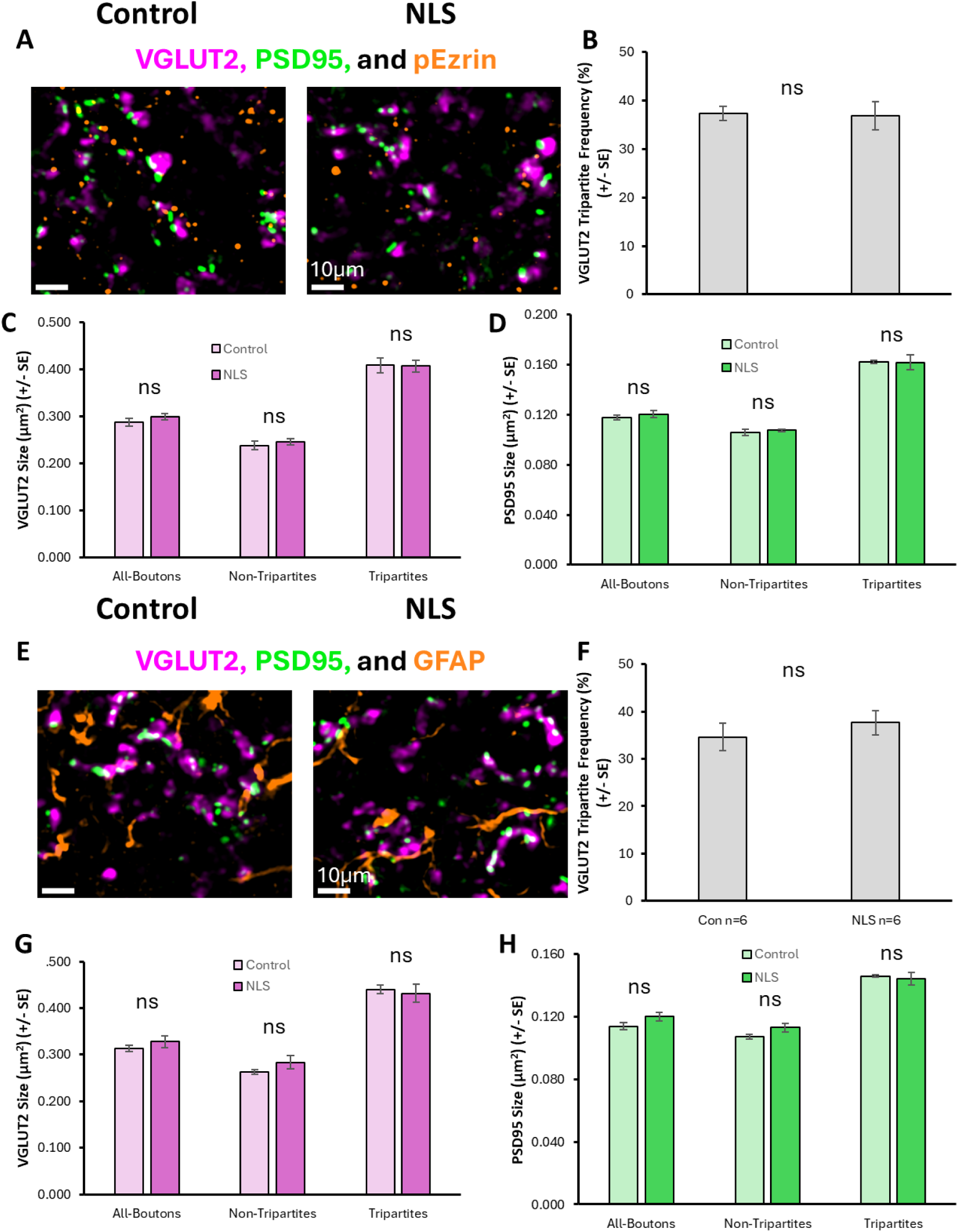
Excitatory VGLUT2-associated Tripartite Synapses in sALS. (A) Immunohistochemical stain of VGLUT2, PSD95 and pEzrin to identify excitatory tripartite synapses. (B) Bar graph plotting the percentage of VGLUT2 synapses that are tripartite (pEzrin contacted) in control and TDP43ΔNLS (NLS) mice. (C) Bar graph of the size of VGLUT2 boutons associated with all synapses, tripartite and non-tripartite synapses, in control and NLS mice. (D) Bar graph of the size of PSDs associated with all VGLUT2 synapses, tripartite and non-tripartite synapses in control and NLS mice. (E) Immunohistochemical stain of VGLUT2, PSD95 and GFAP to identify excitatory tripartite synapses. (F) Bar graph plotting the percentage of VGLUT2 synapses that are tripartite (GFAP contacted) in control and TDP43ΔNLS (NLS) mice. (G) Bar graph of the size of VGLUT2 boutons associated with all synapses, tripartite and non-tripartite synapses, in control and NLS mice. (H) Bar graph of the size of PSDs associated with all VGLUT2 synapses, tripartite and non-tripartite synapses in control and NLS mice

Similarly, we observed no change in the percentage of VGLUT2 synapses that were contacted by GFAP-positive structures between TDP43ΔNLS mice and controls (Figure 6E-F). There was also no change in the size of VGLUT2 presynaptic boutons, whether they were part of GFAP-associated tripartite synapses or not (Figure 6G). VGLUT2-associated PSDs were not significantly different between TDP43ΔNLS mice and controls, whether or not they were associated with tripartite synapses (t(10)=0.378, p=0.714) or non-tripartite synapses (t(10)=1.936, p=0.082) (Figure 6H).

Overall, our findings reveal selective changes in postsynaptic structures of VGLUT1-associated synapses, but not VGLUT2-associated synapses. Our findings also show that, contrary to our hypothesis, there is no selective vulnerability of excitatory tripartite synapses in the TDP43ΔNLS mice.

We next investigated whether astrocytic contacts with cholinergic C-boutons were altered in the TDP43ΔNLS model of sALS. C-boutons were identified as VAChT-positive puncta opposed to SMI-32 labelling, which broadly labelled the soma and processes of motor neurons. VAChT-boutons that were partially colocalised with p-Ezrin puncta were classified as tripartite synapses (Figure 7A).

**Figure 7.**
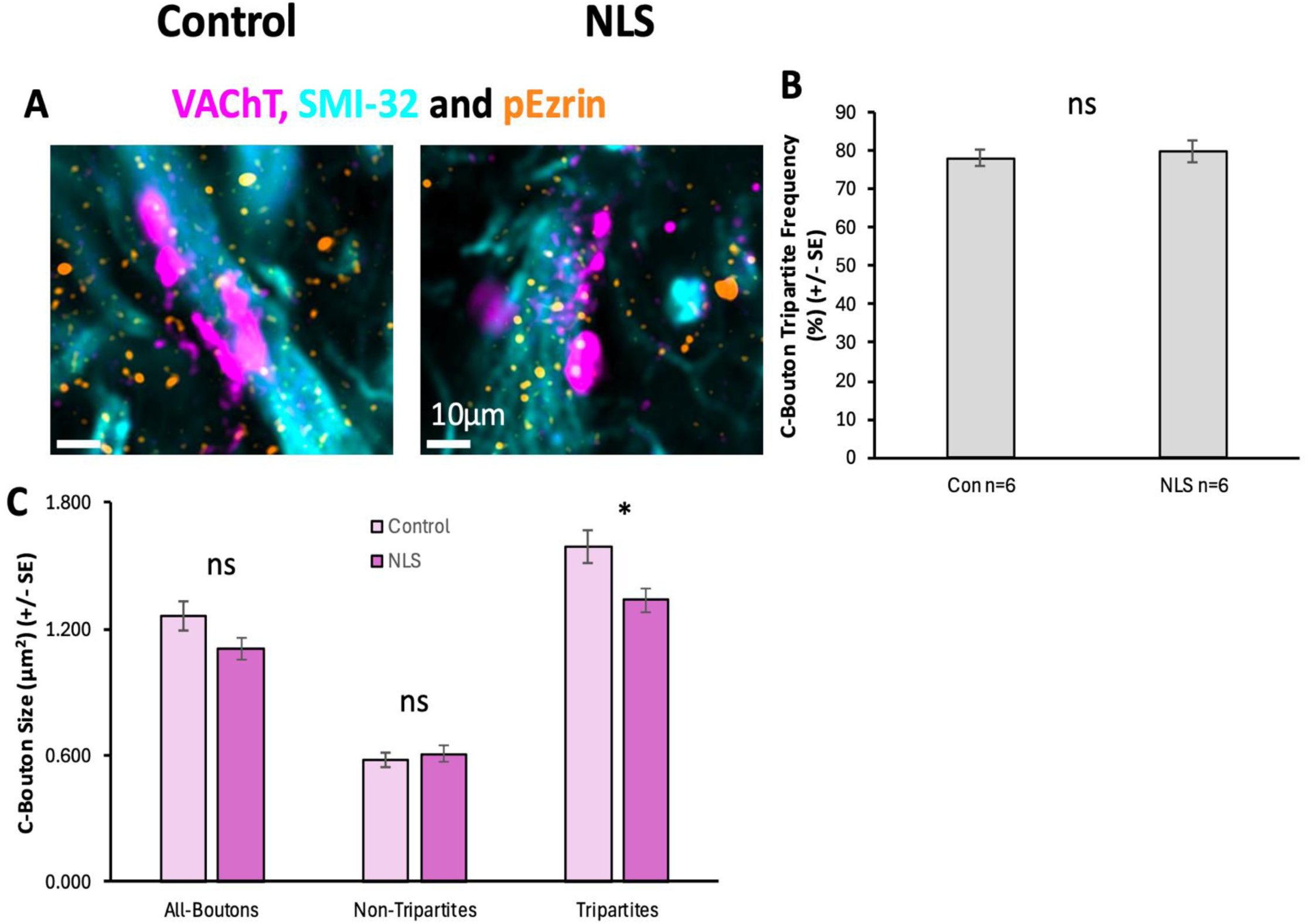
Cholinergic VAChT C-boutons and Tripartite Synapses in sALS. (A) Immunohistochemical stain of VAChT, SMI-32 and pEzrin to identify C-boutons and identify whether they are tripartite synapses or not. (B) Bar graph plotting the percentage of C-boutons that are tripartite (pEzrin contacted) in control and TDP43ΔNLS (NLS) mice. (C) Bar graph of the size of VAChT C-boutons associated with all synapses, tripartite and non-tripartite synapses.

Approximately 80% of all C-boutons were associated with p-Ezrin PAPs (Figure 7B). There was no difference in the percentage of tripartite C-bouton synapses between control and TDP43ΔNLS mice (t(9)=0.469, p=0.650). Interestingly, however, C-boutons associated with p-Ezrin PAPs were significantly smaller in TDP43ΔNLS mice compared to controls (t(9)=2.521, p=0.033). In contrast, there was no significant difference in the size of C-boutons that were not contacted by p-Ezrin PAPs (t(9)=0.596, p=0.566) (Figure 7C). These data may indicate selective morphological changes in tripartite C-bouton synapses.

## Discussion

### C-Boutons in sALS model

Our research demonstrated significant changes in cholinergic C-boutons in the TDP-43ΔNLS mice. C-boutons, which arise from a population of Pitx2-expressing cholinergic interneurons near the central canal of the spinal cord, are distinctively large presynaptic boutons that are associated with a complex post-synaptic organisation including subsurface cisternae ^21,23,42–44^. C-boutons modulate the excitability of motor neurons through activation of muscarinic (m2) receptors and regulation of Kv2.1 channels that leads to reduced spike half-width ^45–47^.

Our observation of overall degeneration of C-boutons replicates prior work in the TDP-43ΔNLS model showing a loss of C-bouton volume and a selective decrease in the number of C-boutons onto fast-type motor neurons innervating the gastrocnemius muscle ^36^. The reduced number of C-boutons in the TDP43ΔNLS model is partly accounted for by the reduced size of motor neurons. When soma size is taken into consideration, gastrocnemius motor neurons in fact display an increased density of C-boutons contacting the soma ^36^. We similarly observed reduced size of MMP9-labelled fast-type motor neurons, which account for approximately half the total number of motor neurons in mid-lumbar ventral horn lateral motor pools ^39,48^.

C-boutons are also smaller in size in the TDP43ΔNLS model. This appears in contrast to the SOD1 model which displays increased C-bouton sizes from early pre-symptomatic stages, which likely acts as a compensatory mechanism for spinal networks ^34,35,49^. One explanation for this difference could an impact of the SOD1 mutation on the formation of synaptic circuits during development, which is not possible in the TDP43ΔNLS model due to its rapid disease progression in adulthood. Alternatively, this could be a specific effect of the TDP-43 pathology observed in this model, highlighting the importance of utilizing models other than the traditional SOD1 model.

Whilst defining the fate of C boutons helps extend our understanding of ALS disease mechanisms, C-boutons also represent potential therapeutic targets due to their modulation of motor neuron function. Experimental silencing of C-boutons has been shown to improve muscle innervation, and combined with task-dependent muscle training, may improve behavioural capabilities of SOD1 animals ^35^, although, our results suggests that this may not necessarily have the same effect in models exhibiting TDP-43 pathology.

### Inhibitory Synapses in sALS model

There is considerable interest in the contribution of different spinal interneuron subtypes to the early stages of ALS, including inhibitory interneurons. However, from analysis of the ubiquitous inhibitory synapse marker, GLYT2, we observe no significant change in inhibitory synapses. Recently, we hypothesised that early developmental changes in excitatory and inhibitory synapses in ALS could pre-dispose spinal motor circuits to an excitatory/inhibitory imbalance, leading to hyperexcitability and excitotoxicity ^32^. Our findings from the SOD1 and C9orf72^50^ mouse models showed there was no such early-stage imbalance in the number of excitatory and inhibitory synapses in spinal neurons both in vitro and in vivo in the first 30 postnatal days ^51^.

However, other studies have demonstrated a reduced number of inhibitory synapses in the brain and spinal cord ^52–54^. Furthermore, studies in the SOD1 mouse model at slightly later, albeit early or presymptomatic, stages (30-60 days) demonstrate both functional and structural changes in inhibitory synaptic input to motor neurons ^33,55,56^. Spatial transcriptomics has shown a significant loss of V1-inhibitory interneurons that precedes the loss of motor neurons ^31^, whilst genetic stabilisation of a vulnerable V1 inhibitory interneuron subpopulation ameliorates the motor phenotype in the SOD1 model ^56^. Functional analysis of recurrent inhibition to motor neurons found that it was significantly reduced in juvenile SOD1 animals, which correlated with reduced density of glycine receptor clusters ^33^. Interestingly, while these changes were observed in young juvenile animals (14-25 days old), in adulthood (40-60 days old) they were comparable to healthy controls suggesting compensatory mechanisms.

Given the molecular and anatomical diversity of inhibitory synapse subtyps, it is possible that analysis using a pan-inhbitory synapse marker such as GLYT2 may occlude subtype-specific changes. It is also possible, that the sporadic TDP43ΔNLS mouse model used in our study, could have shown early changes in inhibitory synapses to motor neurons in the first weeks following induction that are then compensated for by 3-4 weeks post-induction.

### Excitatory Synapse Changes in sALS model

Our results highlight a selective vulnerability of VGLUT1-associated excitatory synapses, whilst VGLUT2-associated synapses were unchanged in their size and number. Within the spinal cord, the majority of VGLUT2 boutons originate from local spinal neurons, with a smaller number arising from nociceptive inputs ^25,57^ and descending inputs from the rubrospinal and vestibulospinal tracts ^58^. VGLUT1 primarily labels Ia afferents from dorsal root ganglia sensory neurons to provide excitatory proprioceptive feedback to large alpha MNs that are most vulnerable to degeneration in ALS ^48,59–61^ ^25,62^.

Our findings demonstrate a loss of VGLUT1 synapses and a reduced size of PSD-95 clusters associated with VGLUT1-boutons. In our previous study, which examined synaptic pTDP-43 clusters in mouse spinal cord tissue, we similarly demonstrated that VGLUT1-associated synapses are more substantially affected in the SOD1 model compared to VGLUT2-associated synapses ^16^. Other studies have shown functional and structural changes in VGLUT1-positive Ia afferents to motor neurons, suggesting a time-course of early-stage increases in the strength of proprioceptive inputs, followed by reduced strength, reduced postsynaptic clustering, and finally a loss of presynaptic VGLUT1 inputs to motor neurons ^16,29^. Spinal muscular atrophy similarly shows early functional and structural impairment in Ia afferent connections to MNs ^63,64^. Recent attempts using trans-spinal direct current stimulation have been successful in restoring Ia afferent structure and function in SOD1 animals, although there was no change in the disease burden in the animals, as determined by the expression of phosphorylated pCREB ^65^.

The concept of a conserved VGLUT1-associated synaptic phenotype leading to comparable motor deficits in two mouse models (i.e. SOD1 and TDP43ΔNLS) with fundamentally distinct genetic and pathological mechanisms, is of particular interest for understanding the molecular mechanisms of ALS. This may implicate other shared downstream targets of both SOD1 and TDP-43. Indeed, there is conflicting evidence on the role of TDP-43 in ALS-synaptopathy. TDP-43 is present at synapses ^16^, it is implicated in the translation of synaptic proteins ^13,17^, and mutations in the gene encoding TDP-43 (TARDBP) lead to altered dendritic arborisation, spine formation and synapse numbers in cortical neurons and corticospinal circuits ^30,66,67^. Synaptic analysis from human ALS post-mortem brain tissue shows a correlation between patient TDP-43 pathology and the degree of synapse loss ^68^. However, in the spinal cord, synapse loss does not correlate with the number of neurons showing TDP-43 pathology ^69^.

### No Astrocyte Reactivity to Neuronal TDP-43 pathology

In this model the TDP-43 is exclusively driven in neurones using a neurofilament heavy polypeptide promoter. This allows us to determine the neuron autonomous effects of the pathology. This in important as there is considerable evidence of non-cell autonomous mechanisms either driving or exacerbating pathology in ALS ^70–72^. Molecular studies implicate changes in neural-glial signalling mechanisms in the spinal cord of the SOD1 mouse model ^73^. Histologically a selective loss of tripartite synapses across ventral and intermediate laminae of the lumbar spinal cord is observed in the SOD1 mouse model and in post-mortem tissue from ALS patients. Spinal cord tripartite synapses are structurally and molecularly enriched in PSD95, which may in part be driven by astrocyte activity ^24^. Tripartite synapses harbour the molecular machinery for a bi-directional neural-glial signalling mechanism whereby astrocytes provide modulatory feedback inhibition of spinal motor networks that may act to prevent hyperexcitability ^74^. Many individual components of this neural-glial signalling pathway, such as mGluR5 signalling and purinergic signalling have been independently verified as dysfunctional or deficient in ALS ^75–78^.

Given this body of information, we hypothesised that tripartite synaptopathy is a conserved hallmark of ALS. However, our findings from this TDP43ΔNLS model suggest that there is no evidence of astrocyte pathology or selective excitatory tripartite synapse loss. For example, the overall fraction of either VGLUT1-associated or VGLUT2-associated excitatory tripartite synapses was unchanged, and structural changes in PSDs was independent of astrocytic contacts. Although C-boutons showed a degree of selective structural changes when associated with p-Ezrin, there was no overall change in the fraction of C-boutons contacted by p-Ezrin. Indeed, unlike previous observations from the SOD1 model, there were no changes in the number of p-Ezrin PAPs in the TDP43ΔNLS spinal cord.

These data support there being no such astrocyte pathology in this model. Therefore, whilst there is interest in the non-cell autonomous mechanisms of ALS, such as astrocytic or microglial cell pathology, some of the synaptic hallmarks of the disease (such as structural changes and synapse loss) may in fact be driven by neuronal mechanisms.

Understanding the differential contribution of TDP-43 pathology in neuronal and glial cell populations, separately and combined, is necessary to dissect the cellular mechanisms of ALS. Future work driving the ΔNLS TDP-43 transgenes selectively in glia will allow us to further address this.

## Conclusion

Whilst there is evidence that synapses are structurally and functionally altered in the early stages of ALS, what is lacking is a comprehensive understanding of which types of synapses are vulnerable. Our investigation begins to tackle this by examining a range of synapse subtypes in the spinal cord, and contributing to evidence of selective synapse vulnerability in a model of sALS. We also observe hallmarks of synaptopathy that are seemingly conserved across models of ALS, indicating different molecular mechanisms underlying ALS may converge on particular synapses and circuits.

## Acknowledgements

The authors thank Dr Carmen Falcone and Dr Simon S. Sharples for their feedback and critical reviews that helped refine the clarity and rigor of the manuscript.

## Author Contributions

AAH designed and performed experiments and data analysis and wrote the manuscript. EB performed experiments and data analysis and contributed to the writing. CFM undertook in vivo experiments on the TDP43ΔNLS mice and processed the tissue for shipment from Copenhagen to St Andrews, as well as contributed to writing the manuscript. GBM helped design the experiments and contributed to the writing of the manuscript. MJB designed and performed experiments and data analysis, contributed to writing the manuscript and supervised the project.

## Data Availability

Data analysis files will be included in a data repository. Raw image data will be made available upon request to the corresponding author.

